# 40 Hz Steady-State Response in Human Auditory Cortex Is Shaped by GABAergic Neuronal Inhibition

**DOI:** 10.1101/2023.10.20.563259

**Authors:** Alessandro Toso, Annika P. Wermuth, Ayelet Arazi, Anke Braun, Tineke Grent-‘t Jong, Peter J. Uhlhaas, Tobias H. Donner

## Abstract

The 40 Hz auditory steady-state response (ASSR), an oscillatory brain response to periodically modulated auditory stimuli, is a promising, non-invasive physiological biomarker for schizophrenia and related neuropsychiatric disorders. The 40 Hz ASSR might be amplified by synaptic interactions in cortical circuits, which are, in turn, disturbed in neuropsychiatric disorders. Here, we tested whether the 40 Hz ASSR in human auditory cortex depends on two key synaptic components of neuronal interactions within cortical circuits: excitation via N-methyl-aspartate glutamate (NMDA) receptors and inhibition via gamma-amino-butyric acid (GABA) receptors. We combined magnetoencephalography (MEG) recordings with placebo-controlled, low-dose pharmacological interventions in the same healthy human participants. All participants exhibited a robust 40 Hz ASSR in auditory cortices, especially in the right hemisphere, under placebo. The GABAA receptor-agonist lorazepam increased the amplitude of the 40 Hz ASSR, while no effect was detectable under the NMDA-blocker memantine. Our findings indicate that the 40 Hz ASSR in auditory cortex involves synaptic (and likely intracortical) inhibition via the GABA-A receptor, thus highlighting its utility as a mechanistic signature of cortical circuit dysfunctions involving GABAergic inhibition.

**Significance statement:** The 40 Hz auditory steady-state response is a candidate non-invasive biomarker for schizophrenia and related neuropsychiatric disorders. Yet, the understanding of the synaptic basis of this neurophysiological signature in humans has remained incomplete. We combined magnetoencephalography (MEG) recordings with placebo-controlled pharmacological interventions in healthy human subjects to test the modulation of the 40 Hz ASSR in auditory cortex by two synaptic components that have been implicated in the generation of neuronal oscillations in cortical microcircuits: glutamate N-methyl-aspartate glutamate (NMDA) receptors and gamma-amino-butyric acid (GABA) -A receptors. Boosting GABAergic transmission, but not blocking NMDA-receptors, increased the amplitude of this ASSR. Thus, GABAergic inhibition modulates 40 Hz steady-state responses in auditory cortex.

## Introduction

An important objective of current translational research is the identification of biomarkers of the neural circuit dysfunctions underlying schizophrenia and related disorders, such as autism spectrum disorders. The auditory steady state response (ASSR), an oscillatory brain response to a periodically modulated auditory stimulus, has recently gained attention as a candidate neurophysiological biomarker of schizophrenia (Grent-’t-Jong et al., 2023). The ASSR can be measured noninvasively via electroencephalography (EEG) or magnetoencephalography (MEG) (Tan, Gross and Uhlhaas, 2015; Thune, Recasens and Uhlhaas, 2016). The ASSR peak frequency occurs at stimulus frequencies around 40 Hz (Galambos, Makeig and Talmachoff, 1981; Pastor et al., 2002), pointing to a resonance property of the stimulus-driven neural circuits. A substantial body of evidence indicates a reduction of the 40 Hz ASSR in patients diagnosed with schizophrenia (SCZ) (Kwon et al., 1999; Tan, Gross and Uhlhaas, 2015; Tada et al., 2021; Onitsuka et al., 2022) as well as in individuals at risk of developing the disorder (Grent-’t-Jong et al., 2021; Tada et al., 2021) and first-degree relatives of schizophrenia patients (Hong et al., 2004; Rass et al., 2012). A reduction of the 40 Hz ASSR has also been reported in bipolar disorder (Jefsen et al., 2022) and autism (Wilson et al., 2007).

Developing a mechanistic understanding of the 40 Hz ASSR requires delineating the synaptic basis of the 40 Hz ASSR. As a response to an oscillatory stimulus input to the brain, the ASSR may, in principle, simply result from the linear superposition of a sequence of auditory evoked potentials that are faithfully propagated from the inner ear to auditory cortex (Galambos, Makeig and Talmachoff, 1981; Ozdamar, Bohorquez and Ray, 2007; Bohorquez and Ozdamar, 2008; Presacco et al., 2010). However, some evidence indicates that cortical circuits may amplify ASSRs specifically around the 40 Hz range through intrinsic, recurrent interactions (Pastor et al., 2002; Ross, Picton and Pantev, 2002). Stimulus-induced gamma band (40-80 Hz) oscillations result from the reciprocal interaction between excitatory pyramidal cells and inhibitory cells (Donner and Siegel, 2011; Buzsaki and Wang, 2012), with an important role of gamma-amino-butyric acid (GABAergic) inhibitory interneurons (Cardin et al., 2009; Sohal et al., 2009; Veit et al., 2017) and glutamatergic N-methyl-aspartate (NMDA) receptor mediated drive of those interneurons from pyramidal neurons (Carlen et al., 2012; Jadi, Behrens and Sejnowski, 2016).

The goal of the current study was to determine whether NMDA and GABAA receptors are also involved in the generation of 40 Hz ASSRs. To this end, we performed MEG recordings of 40 Hz ASSRs under placebo-controlled pharmacological manipulation of both, GABAA and NMDA receptors in the same participants. We found a marked increase of 40 Hz ASSRs in auditory cortex under GABAA receptor stimulation, but not under NMDA receptor blockade.

## Material and Methods

### Participants

Twenty-three healthy human participants (mean age 28, range 21-40, 9 females) took part in the study after informed consent and introduction to the experimental procedure. The study was approved by the ethics review board of the Hamburg Medical Association responsible for the University Medical Center Hamburg-Eppendorf (UKE). Exclusion criteria, all of which were assessed by self-report, were: history of any neurological or psychiatry disorders, hearing disorder, history of any liver or kidney disease or metabolic impairment, history of any chronic respiratory disease (e.g., asthma), history of hyperthyroidism or hypothyroidism, Pheochromocytoma (present or in history), allergy to medication, known hypersensitivity to memantine or lorazepam, family history of epilepsy (first or second degree relatives), family history of psychiatric disorders (first or second degree relatives), established or potential pregnancy, claustrophobia, implanted medical devices (e.g., pacemaker, insulin pump, aneurysm clip, electrical stimulator for nerves or brain, intra-cardiac lines), any non-removable metal accessories on or inside the body, having impaired temperature sensation and / or increased sensitivity to heat Implants, foreign objects and metal in and around the body that are not MRI compatible, refusal to receive information about accidental findings in structural MR images, Hemophilia, frequent and severe headaches, dizziness, or fainting, regularly taking medication or have taken medication within the past 2 months.

Three participants were excluded from the analyses: one due to excessive MEG artifacts and the other two due to not completing all six recording sessions. Thus, we report the results from n = 20 participants (7 females).

Participants were remunerated with 15 Euros for the behavioral training session, 100 Euros for each MEG session, 150 Euros for completing all six sessions, and a variable bonus, the amount of which depended on task performance across all three sessions. The maximum bonus was 150 Euros.

### Experimental design

Each participant completed six experimental sessions, each starting with the oral intake of a pill (see below). During MEG, participants were seated on a chair inside a magnetically shielded chamber. Each MEG session was around 2.5 hours in duration and consisted of different tasks including the ASSR task described below. The ASSR task took place usually around 90 min after start of the MEG recording (i.e., 4 h after drug intake).

### Pharmacological intervention

In each of the six experimental sessions, we administered one of two drugs or a placebo (double-blind, randomized, crossover design): the NMDA receptor antagonist memantine (Johnson and Kotermanski, 2006), the GABAA receptor agonist lorazepam (Kienitz et al., 2022), or a mannitol-aerosil placebo. Both drugs and the placebo were each administered on two (randomly selected) sessions. The dosages were 15 mg for memantine (clinical steady-state dose for adults: 20 mg) and 1 mg for lorazepam (common clinical daily dose between 0.5 and 2 mg). The substances were encapsulated identically to render them visually indistinguishable. Participants received the pill 150 min before the start of MEG recordings. This delay was chosen to jointly maximize plasma concentrations of both drugs during MEG recordings (spanning about 2 h): peak plasma concentrations are reached ∼3 to 8 h after memantine administration (Noetzli and Eap, 2013) and 2 - 3 h after lorazepam administration (Greenblatt et al., 1982). Participants were kept under observation during the waiting period following administration of the pill, and their blood pressure and heart rate was recorded every 15 minutes. The first and senior author are both certified medical doctors, and emergency medical care by the University Medical Center Hamburg-Eppendorf was available on campus at any time. The six sessions were scheduled at least one week apart to allow plasma levels to return to baseline (plasma half-life of memantine: ∼60 to 70 h (Noetzli and Eap, 2013); half-life of lorazepam: ∼13 h (Greenblatt et al., 1982)).

### Stimulus and task

During the ASSR task in the MEG, we presented 100 repetitions of 1000-Hz carrier tones (duration: 2 s) that were amplitude modulated (ripple tones) at 40 Hz (Tan, Gross and Uhlhaas, 2015). In a subset of sessions, the stimulus was amplitude modulated (AM) at 36.75 Hz (n = 33 over 125 sessions), enabling us to establish the stimulus-frequency dependence. Stimuli were presented binaurally through inner ear tubes with an interstimulus interval of on average 2 seconds (jittered between 1.5 and 2.5 s, equal distribution). Ten flat tones (same intensity as ripple tones) were randomly intermixed with the ripple tones. Participants were instructed to fixate a translucent screen (viewing distance: 64 cm) and respond to the flat tones via button press. These trials were not included in the MEG analyses. Auditory stimuli were presented at a fixed pressure level of 70 dB at both ears for all sessions.

### Data acquisition

### MEG

We used a CTF MEG system with 275 axial gradiometer sensors and recorded at 1200 Hz, with a (hardware) anti-aliasing low-pass filter (cutoff: 300 Hz). Recordings took place in a dimly lit magnetically shielded room. We concurrently collected eye-position data with a SR-Research EyeLink 1000 eye-tracker (1000 Hz). We continuously monitored head position by using three fiducial coils. After seating the participant in the MEG chair, we created and stored a template head position. At the beginning of each following session and after each block we guided participants back into this template position. We used Ag/AgCl electrodes to measure ECG and vertical and horizontal EOG.

### Magnetic resonance imaging

Structural T1-weighted magnetization prepared gradient-echo images (TR = 2300 ms, TE = 2.98 ms, FoV = 256 mm, 1 mm slice thickness, TI = 1100 ms, 9° flip angle) with 1 × 1 × 1 mm^3^ voxel resolution were obtained on a 3 T Siemens Magnetom Trio MRI scanner (Siemens Medical Systems, Erlangen, Germany). Fiducials (nasion, left and right intra-aural point) were marked on the MRI.

### Data analysis

MEG data were analyzed with a combination of customized scripts (see associated code, which will be made available upon publication) and the following toolboxes: FieldTrip (Oostenveld et al., 2011) version 20201009 for MATLAB (2021a; The Math-Works, Inc., Natick, MA) as well as MNE (Gramfort et al., 2013) and pymeg for Python (https://github.com/DonnerLab/pymeg) established in previous work from our laboratory (Wilming et al., 2020).

### Behavior

For the quantification of task behavior, we analyzed hit rate (percentage of correctly detected flat-tone targets), false alarm rate (percentage of ripple tones incorrectly reported as target flat-tones) and reaction times.

### Preprocessing

We used an automated algorithm to label artifacts in the continuous time series recorded in each session. Sensor jumps were detected by convolving each sensor with a filter designed to detect large sudden jumps and subsequently by looking for outliers in the filter response. Muscle and environmental artefacts (e.g., cars passing by the vicinity of the MEG room) were detected by filtering each channel in the 100–140 Hz or <1 Hz range, and by detecting outliers that occurred simultaneously in many channels. After removing epochs containing head movements, squid jumps and muscle artifacts, the remaining time-series were then subjected to temporal independent component analysis (infomax algorithm), and components containing blink or heartbeat artifacts were identified manually and removed. We applied a notch filter to remove 50 Hz power line noise and its harmonic (100-150 Hz) and a high-pass linear phase FIR filter (order 3) with cut-off frequency of 0.1 Hz. The resulting data were segmented in trial epochs of 3 s duration (0.5 s baseline) time-locked to sound onset and all epochs that contained artifacts as defined above were discarded. Finally, the preprocessed data were down sampled to a sampling rate of 400 Hz. For subsequent sensor level analyses, the MEG data were first submitted to a planar-gradient transformation (Bastiaansen and Knosche, 2000).

### Spectral analysis

The preprocessed data from each sensor were then submitted to spectral analysis using sliding-window Fourier transform (Welch’s method, Hanning tapered, padding: 4 seconds) with a frequency resolution of 0.25 Hz, window length of 500 ms, and step size of 25 ms. For sensor-level analyses, spectral power was then computed, for each time and frequency bin, by taking the absolute value of the (complex-valued) Fourier coefficients and squaring. The ASSR was evaluated by normalizing the power at each time and frequency bin (for topographies and cortical maps: time-averaged power at the stimulus frequency +/-1Hz) with the power in the pre-stimulus baseline interval (averaged across trials and time in the interval -250 to 0 ms before stimulus onset). The stimulus-phase-locked component of the ASSR was evaluated by first averaging the broadband MEG signal across trials in the time domain, followed by spectral analysis and baseline normalization as described above (Donner and Siegel, 2011). Inter-trial phase coherence (ITPC) was computed for each frequency (f) and time window (t) as follows:

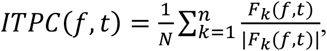

where F denotes Fourier transform, k the trial number and || the complex norm. ITPC ranges from 0 (random phases across trials) to 1 (perfect phase alignment across trials).

### Source reconstruction

We used linearly constrained minimum variance (LCMV) beamforming (Van Veen et al., 1997) to project the MEG data into source space. To this end, we constructed individual three-layer head models (inner skull, outer skull, and skin) from each individual subject’s structural MRI scans. These head models were aligned to the MEG data by a transformation matrix that aligned the average fiducial coil position in the MEG data and the corresponding locations in each head model. Transformation matrices were then generated. We computed one transformation matrix per recording session. We then reconstructed individual cortical surfaces from the structural MRIs and aligned the Glasser atlas (Glasser et al., 2016) to each surface. Based on the head model, we generated a forward model (“leadfields”) for the computation of LCMV filters that was confined to the cortical sheet (4096 vertices per hemisphere, recursively subdivided octahedron). To compute the LCMV filter for each vertex, the leadfield matrix for that vertex was combined with the trial-averaged covariance matrix of the (cleaned and epoched) data estimated for the stimulus interval (0 to 2 s from stimulus onset). We chose the source orientation with maximum output source power at each cortical location. We projected the broadband time-series into source space, and then extracted power (taking absolute value and squaring) for each frequency and time bin. The power values were extracted at each vertex and finally averaged across all vertices within a given region of interest (see next section).

### Regions of interest (ROIs)

We used an established anatomical parcellation of the human cortical surface to define ROIs for further analyses (Glasser et al., 2016). For computing cortex-wide maps of the ASSR, the baseline-normalized power response at the stimulus frequency (+/-1Hz) for the interval from 0.1 to 2 s from stimulus onset was estimated for all 360 parcels of the above atlas. All other analyses collapsed across, the bilateral “early auditory cortex” group, which consists of the following five areas per hemisphere (Glasser et al., 2016): A1, Lateral Belt (LBelt), Medial Belt (MBelt), Para-Belt (PBelt) and retro-insular cortex (RI)). We focused our analyses on this area group as our primary, a priori ROI. We refer to this collapsed ROI as “early auditory cortex” in the following. We used the bilateral central dorsolateral prefrontal cortex group (dlPFC; comprising areas: 9-46d, 46, a9-46v, and p9-46v) as control for anatomical specificity.

### Statistical analyses

Sensors showing a significant 40 Hz ASSR were determined using cluster-based permutation tests (a = 0.05, two-sided, n = 10000) across all 40 Hz sessions under placebo condition, with baseline (−250 to 0 ms) and stimulus-induced (100 to 2000 ms) ASSR power averaged across 39-41 Hz as paired samples. Main effects of stimulation at source-level were determined from source reconstructed ASSR power modulations (expressed in percentage relative to pre-stimulus baseline power) at stimulus frequency +/-1Hz from all 360 Glasser atlas ROIs, using permutation test (two-sided, false discovery rate [FDR]-corrected) across all 20 subjects and sessions under placebo condition. Source-level ASSRs from early auditory cortex (both total and phase-locked power) were also subjected to cluster-based permutation tests across participants (a = 0.05, two-sided, n = 10000) to evaluate if drug differences in MEG power or ITC were significantly different from zero) between 0-2250 ms and stimulus frequency +/-5 Hz (Figure 3A) and ii) between 0-2250 ms averaged across stimulus frequency +/-1 Hz (Figure 3B). Permutation tests were used to evaluate drug differences for the transient component (100-400 ms form stimulus onset and at. stimulus frequency +/-1 Hz) and the sustained component (650-1750 ms from stimulus onset and at stimulus frequency +/-1 Hz) (Figure 3C, Extended Data Figure 3-2). The sustained response interval was chosen so as to avoid any overlap of spectral estimation time windows with the transient or post-stimulus intervals.

## Results

We recorded MEG responses in 20 healthy participants after administration of the NMDA receptor antagonist memantine, the GABAA receptor agonist lorazepam, or a glucose placebo (Fig. 1A; see Methods). Both drugs were applied at relatively low dosage (see Materials and Methods). The ASSR stimuli consisted of 1000-Hz carrier tones (duration: 2 s) that were amplitude modulated at either 40 Hz or 36.75 Hz and presented binaurally (Figure 1B) (Grent-’t-Jong et al., 2021).

**Figure 1.**
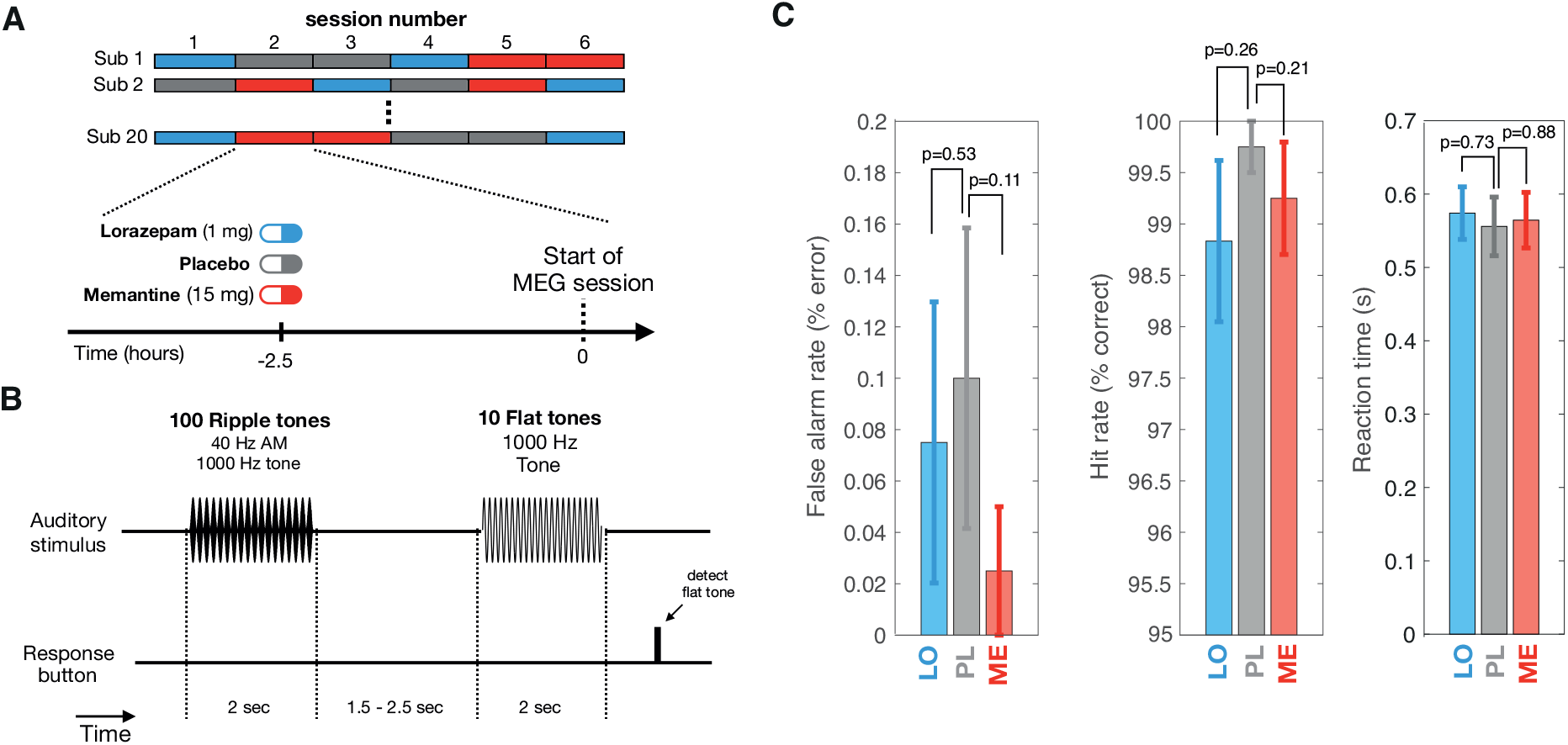
Experimental design, task, and behavior. (A) Experimental design. Subjects underwent six MEG sessions following intake of placebo, lorazepam or memantine and a 2.5 h waiting period. See Materials and Methods for details. (B) ASSR task. Participants had to detect ten 1000 Hz flat tones interleaved in between 1000 Hz tones amplitude-modulated at 40 Hz (ripple tones). (C) Behavioral performance for drug conditions. Error bars, SEM. P-values obtained from two-sided permutation tests (N=1000 repetitions) of differences between drug and placebo conditions.

Participants were instructed to fixate a translucent screen and to detect and respond to flat tones (ten in total) that occurred at random times in between the ripple tones with equal intensity levels over time via button press. The flat tones were presented to control attention but were not used for analyses of the MEG data. The drugs did not affect behavior in this task (Figure 1C, see Materials and Methods), with only a small fraction of participants showing a slightly higher false alarm rate and lower hit rate (6 over 20) compared to maximum performance. Thus, differences in effort or general alertness were unlikely causes of any effects on cortical responses reported below.

### Spatially- and frequency-specific ASSRs

The amplitude-modulated tones produced a robust and specific 40 Hz ASSR during placebo (Figure 2). This response was evident in sensors overlying left and right temporal cortex, with a sustained and significant (p < 0.01, cluster-based permutation test) increase in 40 Hz power relative to pre-stimulus interval over 198 right and left frontal-temporal sensors (Figure 2A). The 40 Hz ASSR was also evident in source reconstructions centered on bilateral auditory cortices), but also including several further temporal, and more distant cortical areas (Figure 2B). The 40 Hz ASSR was stronger in the right hemisphere (Extended Data Figure 2-1 A-B), in line with previous work (Ross, Herdman and Pantev, 2005; Grent-’t-Jong et al., 2021). All our further analyses focused on bilateral early auditory cortex because this is the first stage of auditory cortical processing and it should (as expected), the most prominent ASSRs. The ASSRs detected in neighboring areas may, at least in part, be due leakage from the auditor cortex ASSR through the spatial filters used for source reconstruction (Materials and Methods).

**Figure 2.**
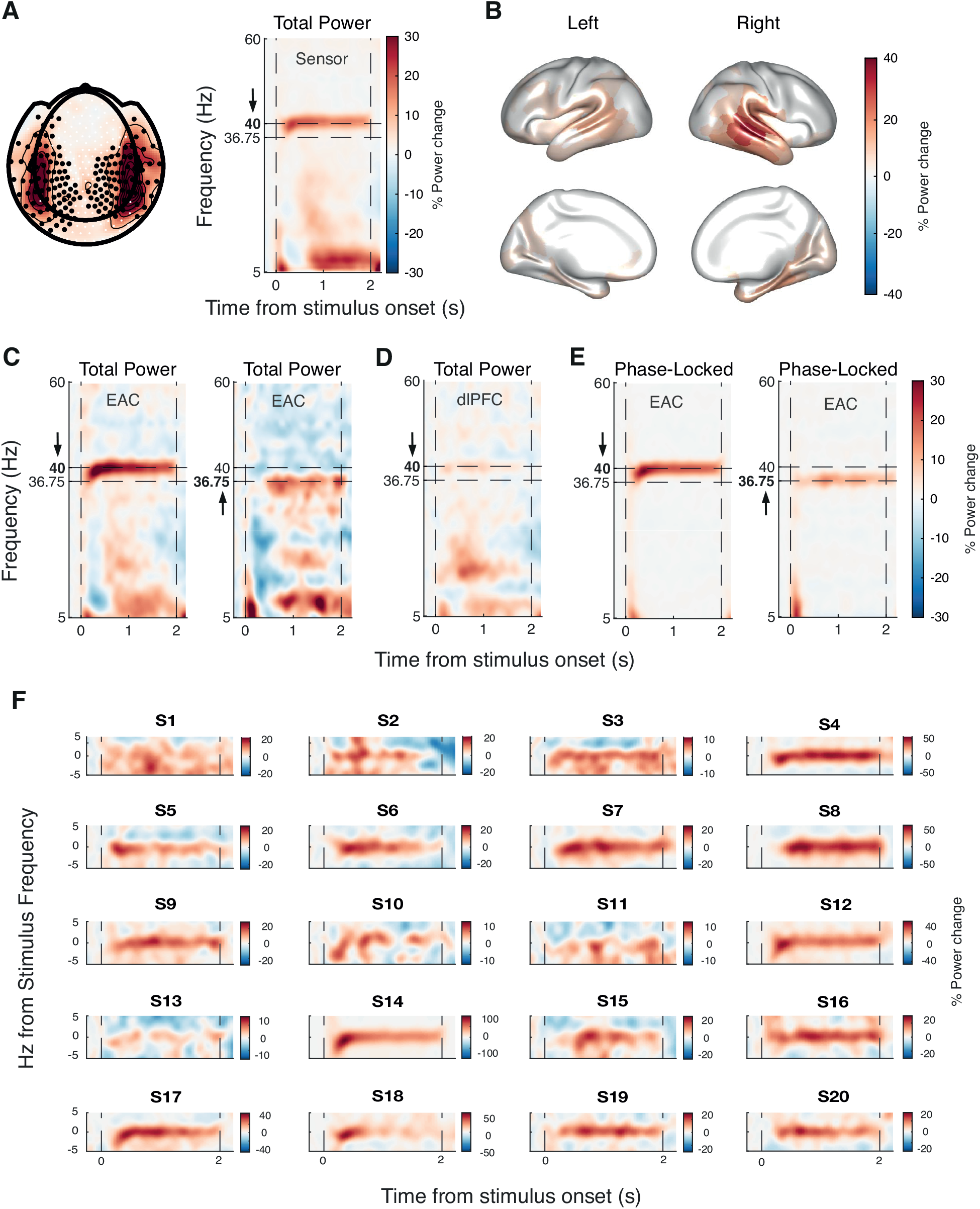
Main effects of ASSR stimulation under placebo condition. **(A)** Left panel depicts % power change relative to pre-stimulus interval at sensor-level averaged across 39 to 41 Hz and 100 to 2000 ms after stimulus onset. Significantly entrained sensors (p < 0.01) are marked with black dots; Right panel shows time-frequency representation of average power change relative to pre-stimulus interval for significantly entrained sensors. **(B)** Significant increases of 40-Hz power across 360 Glasser atlas ROIs. **(C)** Time-frequency representation of power changes relative to pre-stimulus interval in 40 Hz sessions (left panel) and 36.75 Hz sessions (right panel) in early auditory cortex. **(D)** Same as panel C but for dlPFC. (E) Time-frequency representation of phase-locked signal components in 40 Hz sessions (left panel) and 36.75 Hz sessions (right panel) in EAC. (**F)** As panel C, but relative to stimulus frequency and shown separately for each individual (S, subject)

**Figure 3.**
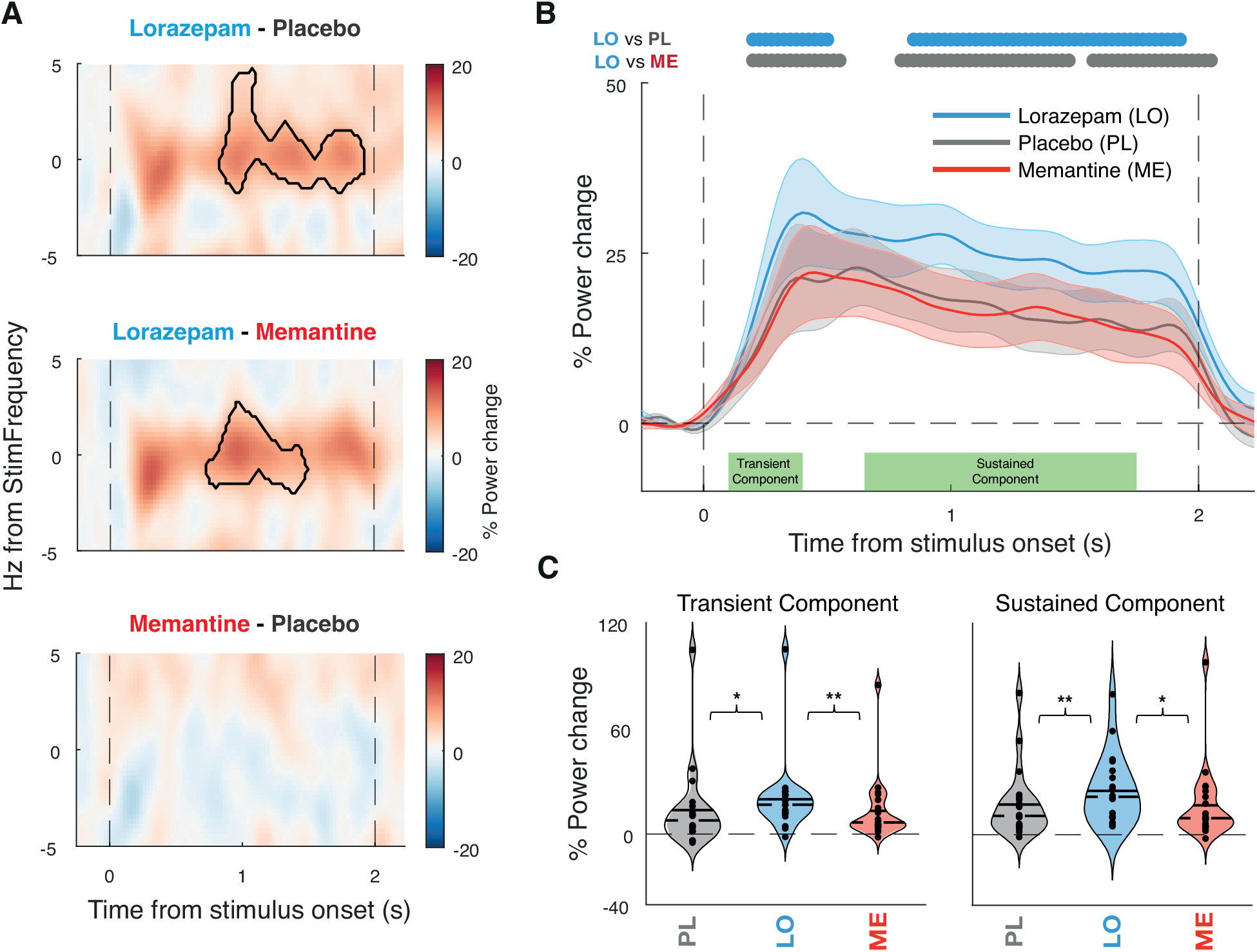
Main drug differences in the auditory steady-state response (ASSR) signals. (A) Time-frequency representation of the power response in early auditory cortex shown as percentage power change between drugs. Contour indicating significant differences between drug conditions (p < 0.05; cluster-based permutation test). Dashed vertical lines indicate stimulus onset and offset. (B) Percentage of power change from baseline over time averaged across +/-1 Hz from stimulus frequency for each drug. Horizontal bars, condition differences (p < 0.05; cluster-based permutation test) and shaded error bars representing SEM. (C) % Power change for each drug for stimulus frequency (+/-1 Hz), separately for transient (100 to 400 ms from stimulus onset, left) and sustained (650 to 1750 ms from stimulus, right) response components. The sustained response interval was chosen so as to avoid any overlap of spectral estimation time windows with the transient or post-stimulus intervals. Solid line representing the mean and dashed line representing median. Significance estimated via permutation dependent-sample test indicated by stars (* p < 0.05, ** p < 0.01).

When we varied the frequency of the stimulus amplitude-modulation (henceforth “stimulus frequency” for brevity) between 40 Hz and 36.75 Hz, the auditory cortex ASSR was strong in both conditions, with a peak frequency that shifted correspondingly in a dependable manner (Figure 2C, see Extended Data Figure 2-1 C-D for single subject resolution). By contrast, there was no detectable response in other control regions such as the dorsolateral prefrontal cortex (Figure 2D). When isolating the phase-locked component of the stimulus response (Methods and (Donner and Siegel, 2011), we found a similarly sustained and frequency-specific, phase-locked component in the early auditory cortex (Figure 2E). Robust ASSRs were detected within each subject (Figure 2F).

In sum, our stimulus protocol and data analysis approach yielded robust, sustained, and lawful ASSRs in early auditory cortex, with a clear phase-locked component. We next quantified the differences in this ASSR between drug and placebo conditions to assess the involvement of glutamatergic and GABAergic synapses in its generation.

### Boosting GABAergic transmission increases the ASSRs in auditory cortex

Lorazepam increased the 40 Hz ASSR in early auditory cortex relative to both, placebo and memantine, with no detectable effect for memantine versus placebo (Figure 3A, B). Both the early transient response component (100 to 400 ms from stimulus onset, p = 0.015 vs PL, p = 0.004 vs ME) as well as the later sustained component (650-1750 ms from stimulus onset, p = 0.011 vs PL, p = 0.009 vs ME) of the ASSR were increased under lorazepam (Figure 3C). Memantine had no detectable effect on the ASSR. We found no statistically significant effect on the phase-locked response component when assessed in isolation (Extended Data Figure 3-1 A-B), with a significant increase in power under lorazepam only in the early transient, but not the later sustained component of the ASSR (Extended Data Figure 3-1 C). We also found no statistically significant effect on inter-trial phase coherence (ITPC; (Extended Data Figure 3-2).

## Discussion

To illuminate the synaptic and circuit mechanisms underlying the generation of 40 Hz ASSRs, we performed MEG recordings of ASSRs under placebo-controlled manipulation of both GABAA and NMDA receptors in human participants. We found that boosting GABAergic transmission through lorazepam robustly increased the 40 Hz ASSR amplitude in auditory cortex, relative to both placebo and the NMDA receptor antagonist memantine. Our findings are consistent with previous animal work applying other GABAergic drugs and recordings in hippocampus (Vohs et al., 2010; Vohs et al., 2012) or temporal cortex (Yamazaki et al., 2020) and establish a role of GABAergic synaptic transmission in the generation 40 Hz ASSR in human auditory cortex.

Stimulus-evoked, phase-locked responses and response amplification (i.e., resonance) through intrinsic cortical circuit interactions differentially shape the transient and sustained components of cortical power responses (Donner and Siegel, 2011), with early transient response components reflecting predominantly the stimulus-evoked activity and late sustained components reflecting intrinsic circuitry (Ross, Picton and Pantev, 2002; Tada et al., 2021; Grent-’t-Jong et al., 2023). Here, we found that the lorazepam effect was evident in both the early and sustained parts of the total spectral power response at 40 Hz (Fig. 3B, C). By contrast, for the phase-locked component of the 40 Hz, the lorazepam effect was only evident in the early, but not the sustained, part of the response (Extended Data Figure 3-1 B, C) and no effect was found on inter-trial phase coherence (Extended Data Figure 3-2). This is in line with a role of GABAergic circuit interactions in the intrinsic amplification of the ASSR within early auditory cortex. Such recurrent cortical interactions may introduce small phase jitter that leads to cancellation when averaging across trials in order to isolate the phase-locked component (Donner and Siegel, 2011).

Previous work has shown that the impact of NMDA receptor blockade on 40 Hz ASSRs may vary dependent on the type of open channel blocker, dose, or the mode of drug administration. Subcutaneous administration of the NMDA-R antagonist ketamine in rodents resulted in an increase of ASSRs power (Vohs et al., 2012; Munch et al., 2023). Another rodent study using acute intravenous ketamine infusion yielded, likewise, a 40 Hz ASSR amplitude increase at low dosage, but an amplitude reduction at higher dosage (Sivarao et al., 2016), indicating a non-linear dose-response relationship. Ketamine infusion in humans also yielded a 40 Hz ASSR amplitude (Plourde, Baribeau and Bonhomme, 1997). NMDA receptor blockade via MK-801in anesthetized rats (Sivarao et al., 2013) or its injections confined to the thalamus (Wang et al., 2020), both decreased the 40 Hz ASSR amplitude, while another study reported an increase in 40 Hz ASSR power when MK-801 was administered intraperitoneally in awake rats (Sullivan et al., 2015). One human EEG study administering memantine found a 40 Hz ASSR increase at frontal electrodes at 20 mg dosage, but no effect at 10 mg (Light et al., 2017). The lack of a detectable memantine effect on the 40 Hz ASSR in auditory cortex in our current study may, therefore, be due to the lower dosage, the small sample size, the time from ingestion (short relative to memantine kinetics, see Materials and Methods), or a combination of these factors. Future work focussing on NMDA receptor effects could use longer waiting times and systematically manipulate the dosage to test for possible effects at higher dosages than the one used here.

Optogenetic manipulations of different interneuron types in animals have established the importance of intracortical inhibition via PV+ interneurons (Bartos, Vida and Jonas, 2007; Buzsaki and Wang, 2012) (Sohal et al., 2009) as well as SOM+ interneurons (Veit et al., 2017) in the intrinsic generation of gamma-band oscillations. 40 Hz ASSRs differ from such intrinsically generated gamma-band oscillations in being driven by a temporal modulated stimulus. Yet, it is conceivable, and consistent with our findings, that the inhibitory cortical circuit motifs for the intrinsic generation of gamma-band oscillations are also involved in the resonance in response to an external 40 Hz stimulus. It is however noteworthy that drive of PV+ cortically-projecting interneurons of the basal forebrain can also induce a cortical ASSR (Kim et al., 2015; Hwang et al., 2019), indicating that a GABAergic manipulation may also alter the ASSR via subcortical pathways.

How does a boost of GABAergic synaptic inhibition increase the amplitude of 40 Hz ASSRs in auditory cortex? Neural circuit modeling has highlighted the importance of the decay time and amplitude of inhibitory postsynaptic currents (IPSC) on PV+ interneurons for the generation of 40 Hz ASSRs (Vierling-Claassen et al., 2008). Other modeling indicates a possible key role of basket cells among PV+ interneurons subtypes in modulating the amplitude of the response, likely due to their larger number and their different axonal targets compared to chandelier cells (Metzner, Zurowski and Steuber, 2019). Benzodiazepines such as lorazepam increase both IPSC amplitude and decay time (Pawelzik et al., 1999; Kang-Park, Wilson and Moore, 2004). In these neural circuit models, those two increases would produce opposing effects on ASSR power. The increase in the 40 Hz ASSR under lorazepam observed in our data is consistent with the IPSC amplitude increase, providing new empirical constraints for circuit models of the 40 Hz ASSR generation.

Different lines of evidence point to an impairment of GABAergic cortical inhibition in schizophrenia. Postmortem studies of schizophrenic patients have revealed a reduction in the GABA transporter GAT-1 (Lewis, Hashimoto and Volk, 2005; Konopaske et al., 2006) and a reduction of GAD-67 (Lewis, Hashimoto and Volk, 2005; Straub et al., 2007), an enzyme responsible of GABA synthesis. This downregulation of inhibitory neurons may be secondary to a deficit in the excitatory drive from pyramidal cells to cortical interneurons (Chung, Fish and Lewis, 2016). Indeed, mice lacking NMDA receptors specifically in inhibitory (GABAergic) PV+ interneurons showed a reduced induction of gamma-band oscillations by optogenetic activation, establishing a possible link between NMDA hypofunction, GABAergic down-regulation and cortical gamma oscillations (Carlen et al., 2012). Likewise, magnetic resonance spectroscopic studies of GABA concentrations in humans found individual GABA-levels to correlate to both peak frequency and amplitude of cortical gamma-band oscillations ((Muthukumaraswamy et al., 2009); but see (Cousijn et al., 2014)), and to be reduced in patients diagnosed with schizophrenia (Yoon et al., 2010; Yoon et al., 2020).

One possibility is that the attenuation of 40Hz ASSRs found in many schizophrenic patients (Kwon et al., 1999; Thune, Recasens and Uhlhaas, 2016; Tada et al., 2021; Onitsuka et al., 2022) and people at risk of developing schizophrenia (Grent-’t-Jong et al., 2021; Tada et al., 2021) results from a deficit in inhibitory activity of cortical interneurons. Schizophrenia is associated with aberrant, intrinsically generated gamma-band oscillations (Uhlhaas and Singer, 2010; Bianciardi and Uhlhaas, 2021) and has been explained in terms of an imbalance between cortical excitation and inhibition (Foss-Feig et al., 2017). Both, intrinsic gamma-band oscillations and cortical excitation-inhibition balance depend critically on intracortical, glutamatergic and GABAergic transmission. Our finding of a GABAergic transmission modulation of the 40 Hz ASSRs is consistent with these previous ideas.

In conclusion, our results indicate that synaptic inhibition via the GABA-A receptor plays a key role in the generation of the 40 Hz ASSR in human auditory cortex. Given the established importance of GABAergic inhibition in schizophrenia and other neuropsychiatric disorders, our findings support the utility of 40 Hz ASSR as a non-invasive signature of aberrations of microcircuit-level processes in human cortex.

## Acknowledgments

We thank Karin Reimann for help with subject recruitment, Marlene Petersson, Barbora Schwarzova and Christiane Reissmann for help with data collection, and Christoph Metzner for discussion.

## Author contributions

T.H.D, P.J.U, A.T. and A.B. conceptualized the study. A.T., A.W. and A.A. collected the data. A.T., and A.W. analyzed the data. A.T., A.W., T.H.D., P.J.U., and T.G.J, interpreted the results. A.T., A.W., and T.H.D. wrote the first draft of the manuscript. All authors commented on, and edited, the manuscript.

## Extended Data

**Extended Data Figure 2-1.**
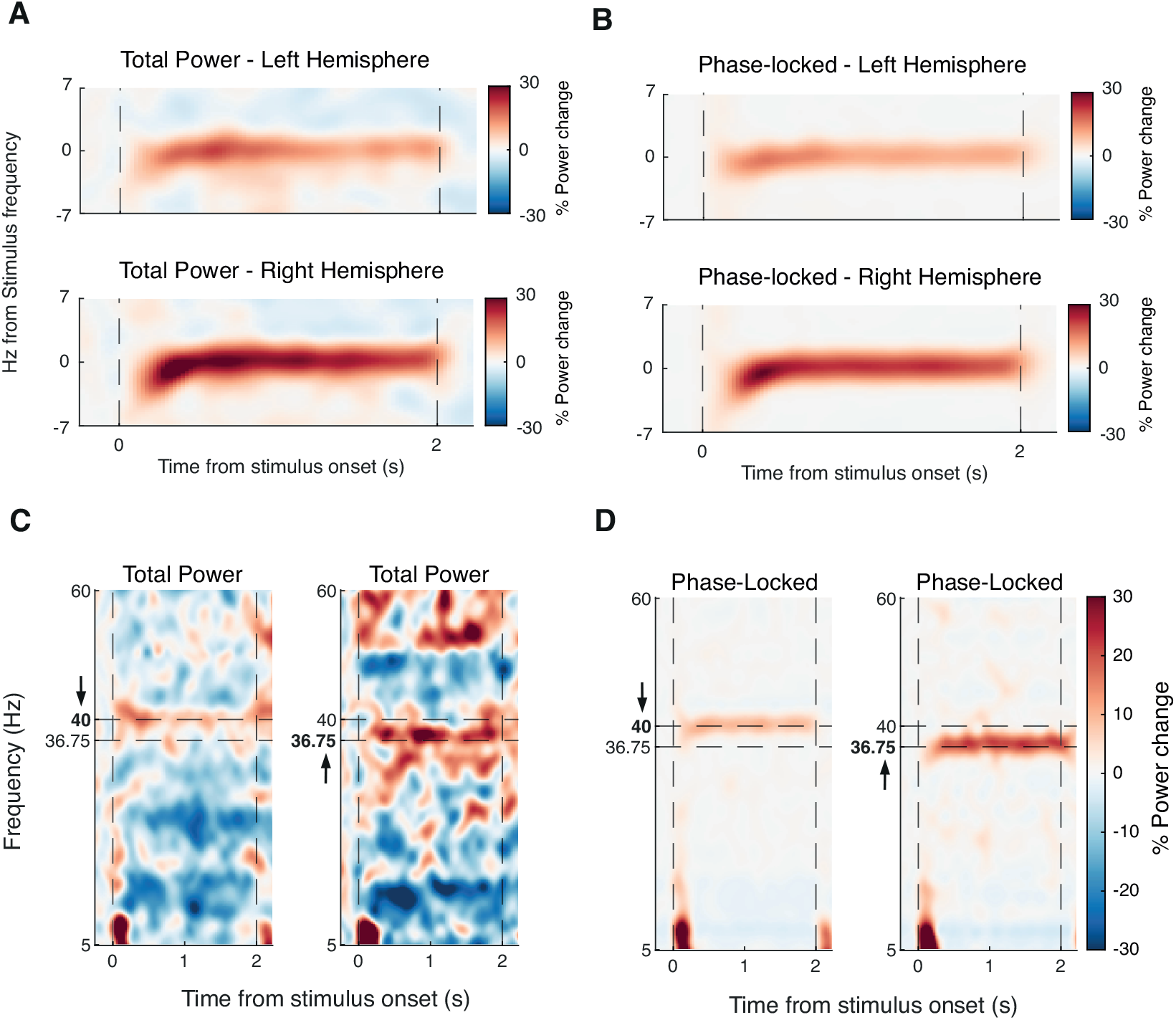
Single subject data and hemispheric lateralization. (A) Time-frequency representation of power changes relative to pre-stimulus interval for left (upper panel) and right (lower panel) EAC. (B) Same as (A) but phase-locked signal components. (C) Time-frequency representation of power changes relative to pre-stimulus interval in a 40 Hz session (left panel) and a 36.75 Hz session (right panel) in early auditory cortex for one subject. (D) Time-frequency representation of phase-locked signal components in a 40 Hz session (left panel) and a 36.75 Hz session (right panel) in early auditory cortex for one subject

**Extended Data Figure 3-1.**
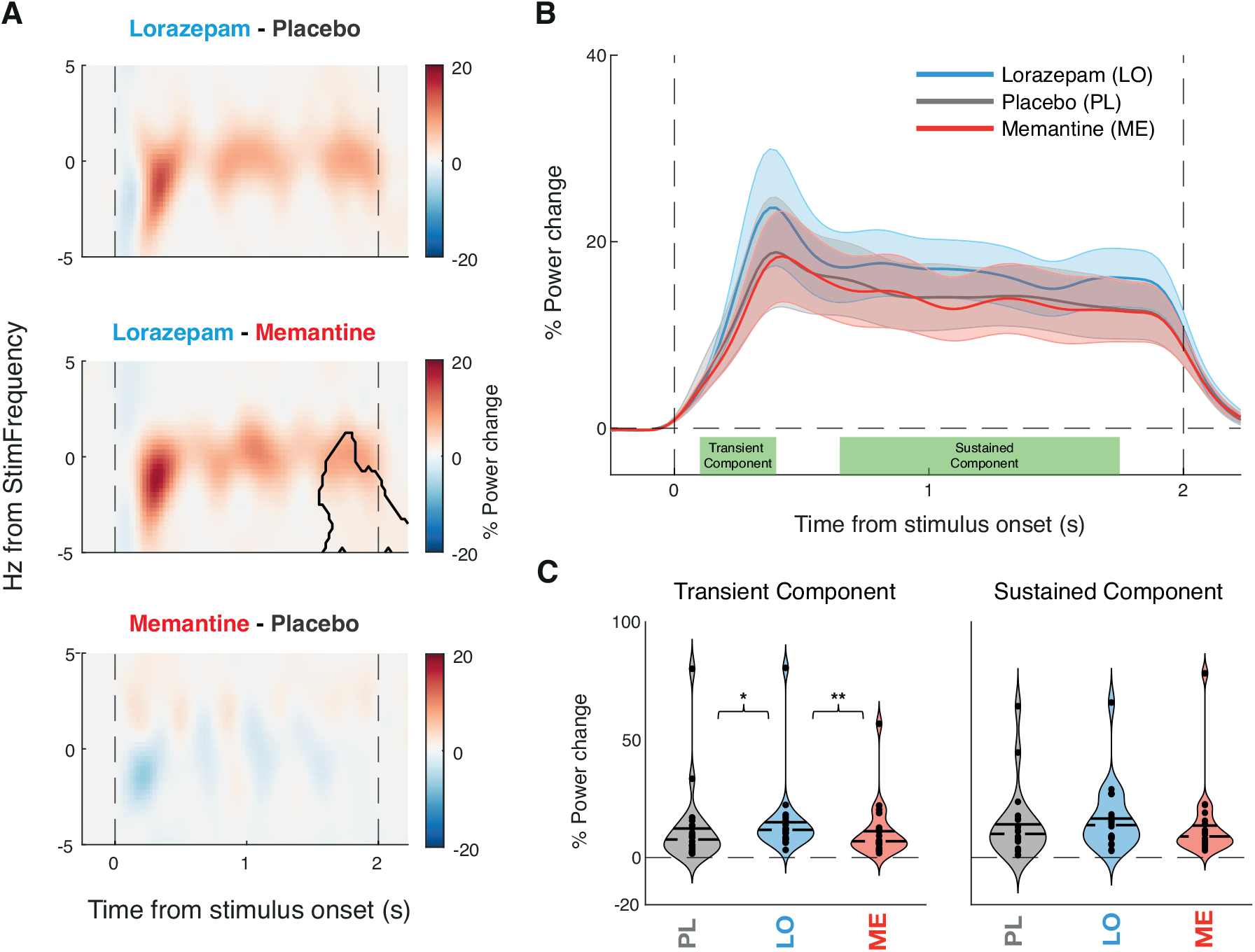
Drug effects on phase-locked response component. (A) Time-frequency representation of the power response in Early Auditory cortex shown as percentage power change between drugs. Contour indicating significant differences between drug conditions estimated via Monte Carlo cluster-based permutation dependent-sample t-test. Dashed vertical lines indicate stimulus onset and offset. (B) % Power change over time averaged across stimulus frequency +/-1 Hz for each drug with dots above indicating significant differences between drugs (p < 0.05) and shaded error bars representing SEM. (C) % Power change for each drug at the stimulus frequency (+/-1 Hz), separately for early (100 to 400 ms, left panel) and late (650 to 1750 ms, right panel) stimulus intervals.. Solid line representing the mean and dashed line representing median. Significance estimated via cluster-based permutation dependent-sample test indicated by stars (* p < 0.05, ** p < 0.01).

**Extended Data Figure 3-2.**
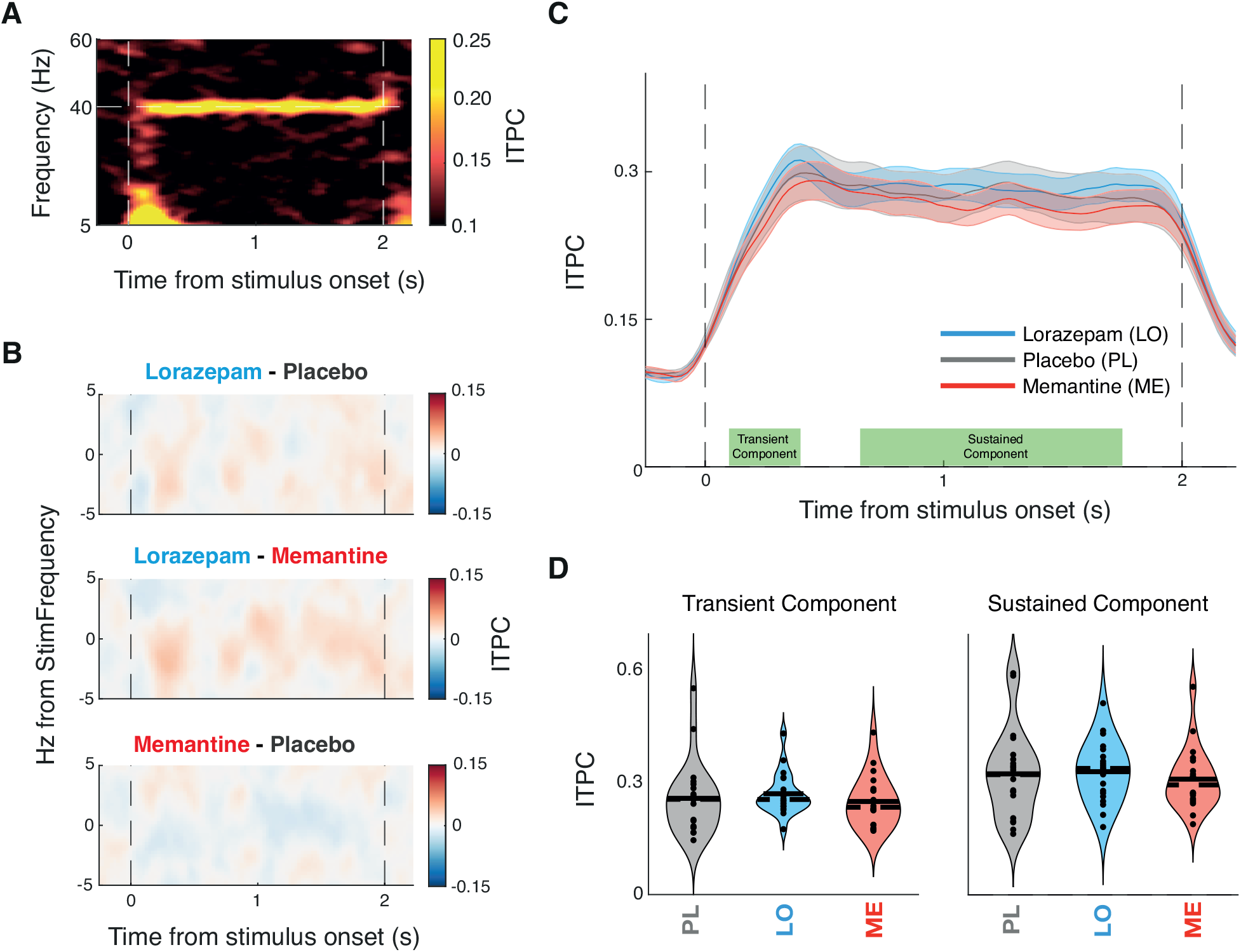
Drug effects on ITPC. **(A)** Time-frequency representation of the ITPC in Early Auditory cortex for placebo sessions in which 40 Hz -modulated stimuli were presented (B) Time-frequency representation of the drug differences in ITPC in Early Auditory Cortex. Contour indicating significant differences between drug conditions estimated via Monte Carlo cluster-based permutation dependent-sample t-test. Dashed vertical lines indicate stimulus onset and offset. (B) ITPC over time averaged across stimulus frequency +/-1 Hz for each drug with dots above indicating significant differences between drugs (p < 0.05) and shaded error bars representing SEM. (C) ITPC for each pharmacological condition at the stimulus frequency (+/-1 Hz), separately for early (100 to 400 ms, left panel) and late (650 to 1750 ms, right panel) stimulus intervals. Solid line representing the mean and dashed line representing median. Significance estimated via cluster-based permutation dependent-sample test indicated by stars (* p < 0.05, ** p < 0.01).

